# YeastSAM: A Deep Learning Model for Accurate Segmentation of Budding Yeast Cells

**DOI:** 10.1101/2025.09.17.676679

**Authors:** Yonghao Zhao, Zhouyuan Zhu, Sen Yang, Weihan Li

## Abstract

An essential step for quantitative image analysis is cell segmentation, which is the process of defining the outline of individual cells in microscopy images. Segmentation of budding yeast is challenging due to their asymmetric cell division and mother-bud morphology. As a result, a dividing cell is frequently misidentified as two separate cells, causing errors in downstream analysis. Here, we overcame this challenge by developing YeastSAM, a deep learning-based segmentation framework derived from µSAM and optimized for budding yeast. YeastSAM achieves more than threefold higher accuracy in segmenting dividing cells compared to existing methods. When combined with single-molecule RNA imaging and organelle imaging, YeastSAM facilitates quantitative analysis of the spatial regulation of gene expression. This study offers an accessible, high-accuracy model for yeast cell segmentation, empowering researchers with minimal programming experience to perform quantitative image analysis.

## Introduction

Imaging methods provide insights into cell morphology and the spatial organization of gene expression [1, 2]. A critical step in image analysis is cell segmentation, which is the computational process of delineating the boundaries of individual cells in microscopy images [3]. When combined with fluorescence imaging, segmentation enables quantitative analysis at the single-cell level, such as variability in gene expression and organelle structure [4, 5]. In recent years, new segmentation approaches have been developed based on convolutional neural networks [6–9] and vision foundation models [10–13]. These methods greatly improve robustness and accuracy compared to traditional image-processing techniques. Frameworks such as µSAM extend these advances by leveraging large-scale pretraining to achieve generalization across different microscopy modalities, such as light and electron microscopy [11].

Despite these advances, segmenting budding yeast *Saccharomyces cerevisiae* remains challenging. During yeast cell division, a bud emerges as a small protrusion from the surface of the mother cell and gradually enlarges until cytokinesis is complete [14, 15]. For most of the cell cycle, the mother and bud remain physically connected [15]. We noticed that current algorithms frequently misclassify the bud and mother as two independent cells, rather than recognizing the mother-bud pair as a single dividing cell. Thus, the morphology of dividing cells often leads to segmentation errors, which in turn compromise downstream analysis. Given the importance of yeast in laboratory research and industrial applications [16, 17], developing an accurate segmentation method is critical. Here, we set out to resolve this challenge.

## Results

### YeastSAM accurately segments yeast cells

Our goal is to develop a segmentation method that accurately identifies budding yeast cells. In this framework, brightfield images are used as input, and the outputs are cell masks (Figure 1A). Brightfield-based segmentation eliminates the need for additional fluorescent markers, preserving fluorescence channels for imaging proteins, RNAs or organelles. This approach also makes the method easier to apply in different experimental contexts. Accurately identifying a budding cell in brightfield images is challenging as it can appear similar to two adjacent G1-phase cells (Figure 1B, C). To address this issue, we performed single-molecule RNA fluorescence in situ hybridization (smFISH) on *CLB2* messenger RNA (mRNA), which encodes a B-type cyclin [18]. Budding cells showed *CLB2* mRNA expression with enrichment in the bud (Figure 1B), which agrees with previous studies [19–21]. In comparison, two adjacent G1-phase cells displayed little *CLB2* mRNA expression (Figure 1C). Thus, *CLB2* mRNA smFISH provides a reference to distinguish budding cells in brightfield images.

**Figure 1.**
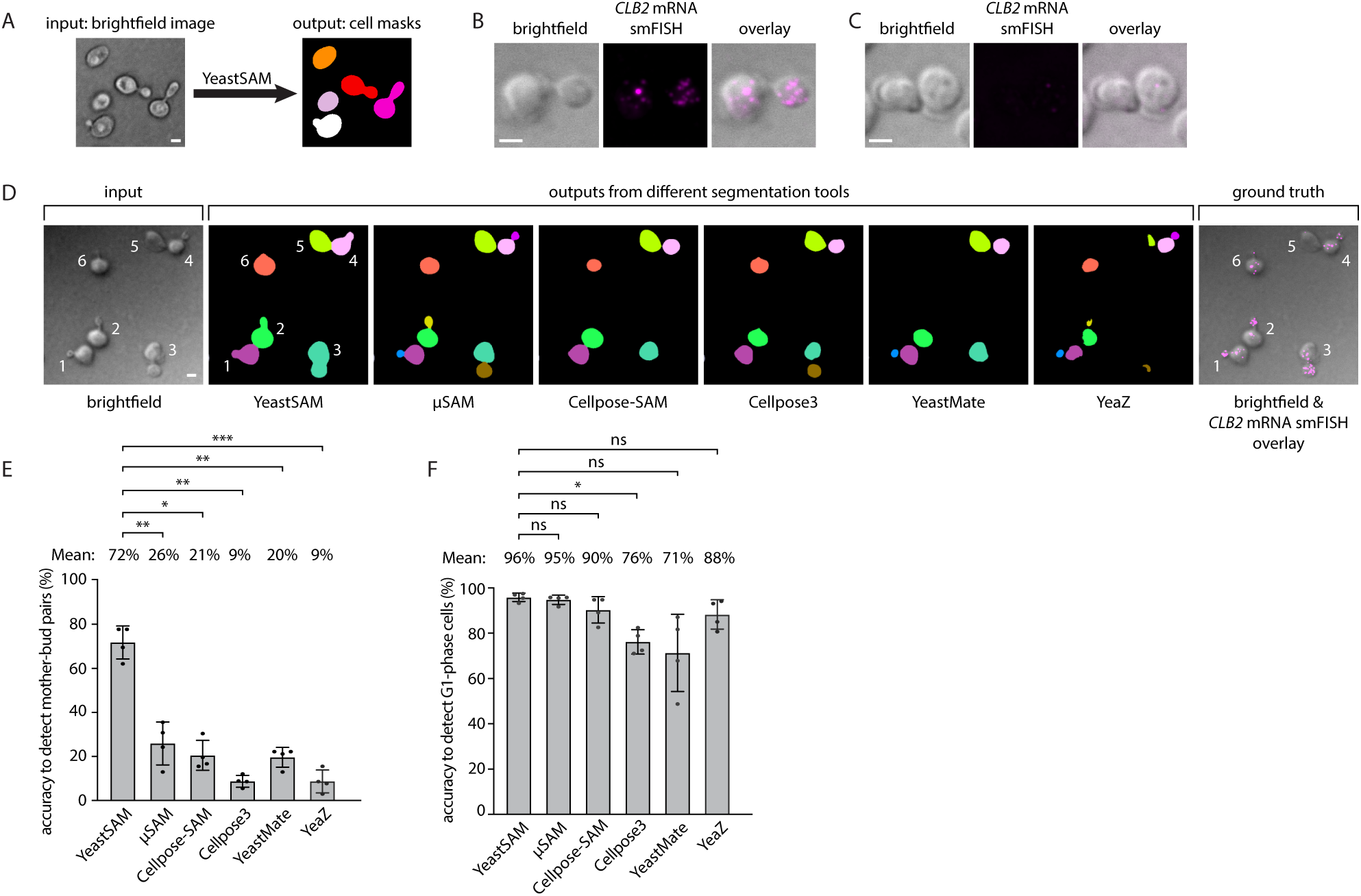
YeastSAM accurately delineates budding yeast cells. (A) Workflow of YeastSAM. Input: 2D brightfield images. Output: cell masks, with different cells shown in distinct colors. (B, C) *CLB2* mRNA smFISH distinguishes a dividing mother-bud pair (B) from two adjacent G1-phase cells (C). Images were acquired as Z-stacks and maximum Z-projected. (D) Comparison of multiple segmentation tools: YeastSAM, µSAM, Cellpose-SAM, Cellpose3, YeastMate and YeaZ. The same brightfield images were used as inputs, and outputs are shown with individual cells in different colors. *CLB2* mRNA smFISH (magenta, rightmost panel) served as a biological reference. A representative field of view with six cells is shown. (E, F) Accuracy of each segmentation method for detecting mother-bud pairs (E) and G1-phase cells (F). Mean values are indicated. P values were calculated using one-way repeated measures ANOVA followed by Tukey’s multiple comparisons test in Prism. Significance is indicated as: *P < 0.05, **P < 0.01, ***P < 0.001, ns: not significant (P > 0.05). Scale bars in Figure 1, 2 μm.

We evaluated the performance of several recently developed segmentation methods, including generalist algorithms such as µSAM [11], Cellpose-SAM [22], and Cellpose3 [23], as well as yeast-specific methods such as YeastMate [6] and YeaZ [24]. Brightfield images served as the program inputs, and the segmentation outputs were manually assessed using *CLB2* mRNA smFISH as a reference. These methods accurately identified G1-phase cells (cell number 5 in Figure 1D) but failed to correctly segment budding cells (cell number 1 to 4 in Figure 1D). Budding cells were misclassified as two separate cells instead of mother-bud pairs. As a result, the accuracy of segmenting budding cells ranged from 9% to 26% (Figure 1E).

To improve segmentation accuracy, we developed a new model. Among the methods we tested, µSAM showed the highest accuracy of 26% in detecting budding cells. Because µSAM is a foundation model for microscopy image segmentation, we reasoned that it could be fine-tuned to create a yeast-specific segmentation framework with improved recognition of mother-bud pairs. To this end, we fine-tuned µSAM on images of 1965 yeast cells that were manually annotated using *CLB2* mRNA smFISH as a reference (Supplementary Movie 1). The resulting fine-tuned model, which we named YeastSAM, generated cell masks that accurately delineate mother-bud pairs (Figure 1D). Notably, YeastSAM achieved 72% accuracy in segmenting budding cells, exceeding the 9-26% achieved by other methods (Figure 1E). For G1-phase cells, YeastSAM reached 96% accuracy, matching or exceeding the performance of existing approaches (Figure 1F). Together, these results showed YeastSAM as a specialized framework for accurate yeast cell segmentation.

### Integrating YeastSAM with FISH-Quant and MitoGraph to analyze mRNA-mitochondria association

Cell segmentation is the first step in quantitative image analysis, providing the basis for downstream measurements of proteins, mRNAs or organelles. As a proof of principle, we applied YeastSAM to analyze mRNA-mitochondria associations (Figure 2A). Previous studies showed that a subset of nuclear-encoded mRNAs is localized to the mitochondrial surface, where their local translation enhances the efficiency of protein import into mitochondria [25–32]. Two well-studied examples are *ATP3* and *TIM50* mRNAs, which encode a subunit of the ATP synthase complex [33] and a component of the Translocase of the Inner Mitochondrial membrane (TIM) complex [34], respectively. Both mRNAs localize to the mitochondrial surface, and disruption of this localization reduces their protein levels, likely due to inefficient import into mitochondria [31]. To examine their localization relative to mitochondria, we imaged the mRNAs using smFISH and mitochondria using a green fluorescent protein (GFP) targeted to the mitochondrial matrix. Both mRNAs localized to mitochondria (Figure 2B, C), consistent with previous reports [31].

**Figure 2.**
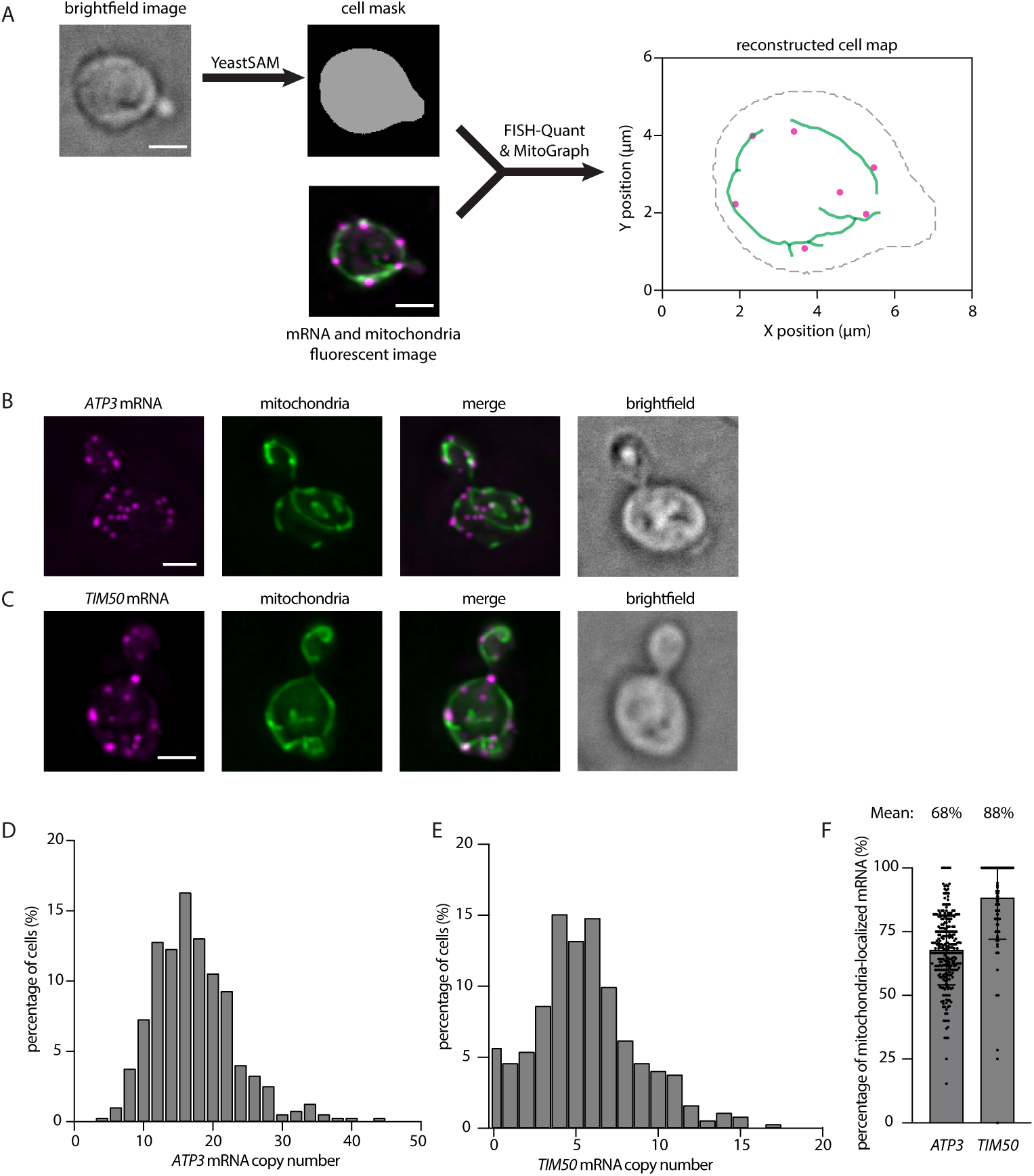
Integration of YeastSAM with FISH-Quant and MitoGraph enables mRNA-mitochondria analysis. (A) Workflow integrating YeastSAM with FISH-Quant and MitoGraph. A representative cell is shown. *TIM50* mRNA smFISH (magenta) and mitochondria labeled with mitochondria-targeted GFP (green) were imaged. Brightfield images were acquired in 2D, while fluorescence images were acquired as Z-stacks and maximum Z-projected. FISH-Quant and MitoGraph detected the XYZ coordinates of individual mRNA molecules and mitochondrial skeletons, respectively. Reconstructed cells combine mRNA positions (magenta), mitochondrial skeletons (green) and cell outlines (gray, XY coordinates). (B, C) Representative images of *ATP3* mRNA (magenta in B), *TIM50* mRNA (magenta in C) and mitochondria (green). mRNAs were detected by smFISH, and mitochondria were labeled with mitochondria-targeted GFP. Images were maximum Z-projected. (D, E) Histograms of *ATP3* (D) and *TIM50* (E) mRNA copy numbers per cell. A total of 391 and 1944 cells were analyzed, respectively. (F) Percentage of mitochondria-localized *ATP3* and *TIM50* mRNAs. Each point represents one cell. mRNAs within 500 nm of mitochondria were classified as mitochondria-localized. Mean values are indicated. Scale bars in Figure 2, 2 μm.

To quantitatively analyze this localization, we established a computational workflow by integrating YeastSAM with FISH-Quant, which detects mRNA positions [35, 36], and MitoGraph, which reconstructs mitochondrial networks [37, 38]. First, YeastSAM generated cell outlines from brightfield images. Next, the mRNA and mitochondrial fluorescence images, along with the cell outlines, were used as the inputs for FISH-Quant and MitoGraph. The final output was a reconstructed cell map containing cell boundaries, mRNAs and mitochondria (Figure 2A). From the reconstructed cell maps, we quantified the number of mRNAs per cell and calculated the fraction of mRNAs that are localized to mitochondria. On average, each cell contained about 15 copies of *ATP3* mRNA and 6 copies of *TIM50* mRNA (Figure 2D, E). The higher abundance of *ATP3* mRNA is consistent with its greater protein abundance compared to *TIM50* [39]. Approximately 68% of *ATP3* mRNAs and 88% of *TIM50* mRNAs were localized to mitochondria, consistent with previous reports showing that *TIM50* mRNA exhibits higher mitochondrial localization than *ATP3* [31] (Figure 2F). These results demonstrate that integrating YeastSAM with FISH-Quant and MitoGraph enables quantitative analysis of mRNA-mitochondria associations.

### Delineating mother and bud boundaries

During yeast cell division, a set of proteins, mRNAs and organelles are asymmetrically partitioned between the mother cell and the bud [40]. This process ensures organelle inheritance in daughter cells [41, 42], controls mating-type switching [43] and regulates replicative aging [44, 45]. While YeastSAM provides accurate cell segmentation, it does not distinguish mother and bud compartments. To address this need, we developed a mother-bud separation module, which integrates with segmentation approaches to delineate mother-bud boundaries. Using the cell outlines generated by YeastSAM as input, this module first distinguishes budding cells from G1-phase cells. Then, it applies a U-Net [46] to draw a boundary line that separates the mother and bud (Figure 3A). By manual inspection, the module achieves an accuracy of 92% and 88% in distinguishing G1-phase cells and budding cells, respectively (Figure 3B), and 91% accuracy in drawing the mother-bud boundary line. As an example, we applied our integrated workflow— combining YeastSAM, the mother-bud separation module and FISH-Quant—to analyze the asymmetric distribution of *CLB2* mRNA [21]. In this workflow, YeastSAM generated cell outlines, the mother-bud separation module defined the mother-bud boundaries, and FISH-Quant detected individual mRNAs. The outputs were reconstructed cell maps, showing cell outlines, mother-bud boundaries and mRNA coordinates (Figure 3A). Using these reconstructed maps, we estimated the volumes of the mother and bud by rotating the 2D cell outlines around their central axis to generate a 3D approximation of each compartment [47] Based on these volumes, we calculated the concentration of *CLB2* mRNA. Its concentration in the bud was 3.5 times higher than in the mother (Figure 3C), consistent with the mRNA’s enrichment in the mother [21]. These results demonstrate that the mother-bud separation module, in combination with YeastSAM, enables quantitative analysis of asymmetric molecular distributions within dividing yeast cells.

**Figure 3.**
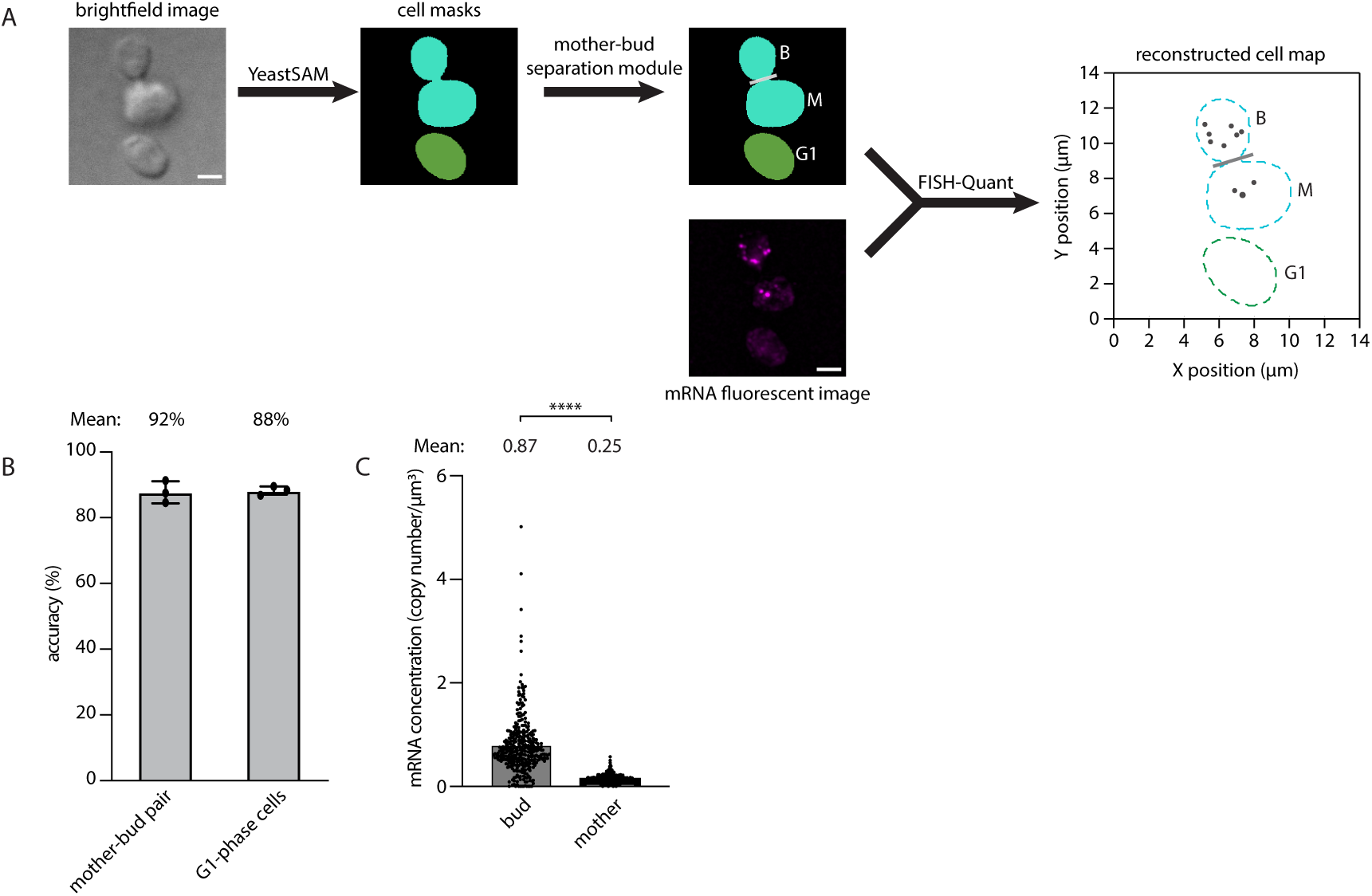
YeastSAM and mother-bud separation module enable analysis of molecular partitioning between mother and bud. (A) Workflow of the integrated approach. *CLB2* mRNA smFISH (magenta) was imaged. Cell masks generated by YeastSAM were used as input for the mother-bud separation module, which distinguishes mother–bud pairs from G1-phase cells. For dividing cells, the module draws a boundary line (grey) that separates the mother and bud and labels them (M: mother; B: bud; G1: G1-phase cell). (B) Performance of the mother-bud separation module in classifying mother-bud pairs and G1-phase cells. Mean values are indicated. (C) Quantification of *CLB2* mRNA concentrations in mother and bud compartments. Each point represents one cell (n = 465). P values were calculated using one-way repeated measures ANOVA followed by Tukey’s multiple comparisons test in Prism. ****P < 0.0001; mean values shown. Scale bars in Figure 3, 2 μm.

## Discussion

In this study, we developed YeastSAM, a segmentation framework tailored for budding yeast. YeastSAM outperformed both generalist algorithms such as µSAM and yeast-specific tools such as YeastMate and YeaZ. The main limitation of previous methods lies in biases toward convex cell shapes, introduced by training data annotation, loss function design and geometric priors. Most segmentation datasets recognize budding events as two convex cells rather than a concave mother–bud pair [6]. Region-based loss functions further prioritize interior coverage over boundary precision, leading to delineation of concave interfaces [48]. In addition, geometric priors such as convex vector flow fields in Cellpose amplify this bias [23, 49]. By contrast, YeastSAM is not constrained by convex priors and was fine-tuned on annotated dividing yeast cells as single concave units. This training enabled YeastSAM to distinguish mother–bud pairs from two adjacent G1-phase cells, possibly through recognizing structures such as the bud neck [47]. Importantly, the training was guided by *CLB2* mRNA smFISH, which provided a biological reference and helped reduce human bias during annotation. Together, these features allowed YeastSAM to accurately capture mother-bud pairs, explaining its higher accuracy compared to existing approaches. YeastSAM is built on the user-friendly napari platform [50], making it accessible to researchers with limited programming experience (Supplementary Movie 1). It should be noted that, while YeastSAM accurately segments dividing yeast cells in brightfield images, its performance on other imaging modalities, such as fluorescent imaging, remains to be validated.

YeastSAM demonstrates how biologically informed annotations can fine-tune a flexible foundation model to accurately segment challenging cell morphologies. This strategy enables precise, quantitative analysis of asymmetric division and complex cellular structures. YeastSAM is freely available on https://yeastsamdoc.readthedocs.io/en/latest/.

## Acknowledgement

The authors thank R. H. Singer, R. Singh, J. Alfonzo, S. Djuranovic, S. Pavlovic-Djuranovic, C. de Graffenried and the members of the Li laboratory for insightful discussions. This work was supported R00GM148788 (W.L and S.Y.).

## Materials and methods

### Yeast strains and plasmids

Strains used in this study were derived from the *Saccharomyces cerevisiae* background W303 (MATa; *ura3-1*; *trp1Δ2*; *leu2-3*,*112*; *his3-11*,*15*; *ADE2*; *can1-100*) [51]. To image mitochondria, GFP fused to the Su9 mitochondrial targeting sequence was inserted at the *HO* locus through homologous recombination and geneticin or kanamycin selection [32]. Cells were grown exponentially in yeast synthetic medium with auxotrophy-complementing amino acids and 2% glucose under continuous shaking at 26 °C.

### Single-molecule fluorescence in situ hybridization

The smFISH was performed as previously described [32], using probes targeting *CLB2*, *ATP3* or *TIM50* mRNAs. Probes were designed and purchased from LGC, Biosearch Technologies. Their sequences are listed in Supplementary Table 1. Briefly, yeast strains were grown overnight at 26 °C in synthetic medium (2% glucose) and auxotrophy-complementing amino acids. Cells were diluted to an optical density of 0.1 at a wavelength of 600 nm (OD_600_ 0.1) in the morning and incubated at 26 °C. After the culture reached OD_600_ 0.2-0.4, cells were fixed by gently shaking at 25 °C for 45 min in 4% paraformaldehyde (32% solution, EM grade; Electron Microscopy Science 15714). Cells were washed with ice-cold buffer B (1.2 M sorbitol and 100 mM potassium phosphate buffer, pH 7.5) and resuspended in 500 μl spheroplast buffer (1.2 M sorbitol, 100 mM potassium phosphate buffer, pH 7.5 and 20 mM ribonucleoside–vanadyl complex (NEB S1402S)). To each sample, Lyticase enzyme (Sigma L2524) was added at 25 units per OD_600_ of cell, e.g. if the OD_600_ was 0.3/ml and 25 ml of cells was needed, 0.3 × 25 units of Lyticase enzyme were added. Cells were then digested for approximately 10 min a 30 °C. When ∼70% of cells were digested, cells were washed and resuspended in 1 ml buffer B and seeded onto 18 mm poly-l-lysine (Sigma)-treated #1.5 (0.16 to 0.19 mm thickness) coverslips (Fisher Scientific). The coverslips were incubated at 4 °C for 30 min to enable cells to attach, washed with buffer B, and stored in 70% ethanol at −20 °C for at least 3 h. The coverslips were washed with 2×SSC at 25 °C twice. A pre-hybridization mix (10% formamide (ACROS organics 205821000) in 2×SSC) was added to the coverslips. During the 30 min 25 °C incubations, the smFISH probes were prepared. For each coverslip, a SpeedVac chamber was used to dry a mixture of 0.125 μl of 25 μM smFISH probe, 2.5 μl of 10 mg μl^−1^ *Escherichia coli* transfer RNA and 2.5 μl of 10 mg μl^−1^ *E. coli* single-stranded DNA. The dried pellet was resuspended in 25 μl hybridization mix (10% formamide, 2×SSC, 1 mg ml^−1^ BSA, 10 mM Ribonucleoside–vanadyl complex (NEB) and 5 mM NaHPO_4_, pH 7.5) and boiled at 95 °C for 2 min. The resuspended smFISH probes were applied to the coverslips for hybridization and incubated in the dark for 3 h at 37 °C. Next, coverslips were washed twice with pre-hybridization mix at 37 °C for 15 min, followed by another three 10 min washes at 25 °C using 2×SSC with 0.1% Triton X-100, 1×SSC, and 1×PBS, respectively. Coverslips were mounted on microscope slides (Thermo Fisher Scientific) using ProLong Gold antifade with DAPI (Thermo Fisher Scientific).

### smFISH and mitochondria image acquisition and analysis

The smFISH images were acquired on an Olympus BX63 wide-field epi-fluorescence microscope using a 100x UPlanApo objective, an X-cite 120 PC lamp (EXFO) and the ORCA-R2 Digital CCD camera (Hamamatsu). Metamorph software (Molecular Devices) was used to control the microscope. Images were acquired at a xy pixel size of 64.5 nm, with Z-stacks acquired at 200 nm step size over an optical depth of 8 μm.

### Data preparation

Our annotations were generated using the µSAM napari plugin . With information from *CLB2* imaging, we can determine which cells are budding and generate ground truth (GT) masks covering each mother-bud pair. In cases where µSAM initially segmented a budding event into separate mother cells and buds, we applied point prompts to merge them into a single complete mask representing the entire budding cell (Supplementary Movie 1).

We developed an additional mask editor for researchers to manually separate the mask of mother and bud cells in each budding event. With this editor, users draw a dividing line between mother and bud and assign cell numbers. By convention, suffix “01” indicates a bud cell, and suffix “02” indicates a mother cell. Cells without suffix are annotated as non-budding. Cell number will be set as the pixel value, we also outputted outline files in a FISH-Quant-compatible format.

Under this annotation rule, we constructed two datasets. The first contained 1,965 cells for training YeastSAM, generated with the µSAM napari plugin and *CLB2* smFISH as reference (brightfield-mask pairs, without mother-bud pair separation). The second contained 810 cells, where our mask editor was used to label each cell as non-budding, mother cell or bud. From this, we derived (i) a budding vs. non-budding classification set and (ii) a mother-bud pair separation set, which were used for evaluation and benchmarking of the mother-bud separation module.

### YeastSAM training

We fine-tuned the default ViT-B-LM, ViT-L-LM and its automatic instance segmentation (AIS) mask decoder of µSAM using three strategies: full fine-tuning (Full FT), freezing the encoder, and low-rank adaptation (LoRA) on the encoder [52]. In the LoRA setting, low-rank matrices *A* ∈ ℝ^*d×r*^ and *B* ∈ ℝ^*r×d*^ (with *r* ≪ *d*) were introduced to approximate the weight updates:

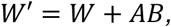

where 𝑊 is the frozen pre-trained weight.

Training was performed with the AdamW optimizer and an initial learning rate of 1e-5. We also tested 1e-4, 5e-5, and 5e-6, and found 1e-5 produced the smoothest convergence. Patch size was set to 512×512, with batch size of 1 and 20 objects per batch on a single RTX 4090 GPU. For LoRA fine-tune, LoRA rank was set to 16 and alpha was set to 32, with a correspondingly enlarged objects per batch of 25. Only matrix Qs and matrix Vs are fine-tuned by LoRA. As an additional augmentation strategy, we applied prompt box perturbation with a box distortion factor of 0.025 to improve robustness. For mask inputs, each instance was processed individually: object masks were transformed into distance maps and resized to the patch size, while the corresponding prompt boxes were converted and perturbed before being passed to the model. The main training follows µSAM’s notebook.

The training loss combined Dice loss with mean squared error (MSE) [53]:

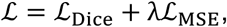

where λ is a balancing weight. Since AIS was enabled, an additional UNETR decoder was trained as implemented in µSAM. This decoder predicts three distance-map channels: foreground probability 𝑓𝑔, distance to object center 𝑐𝑡𝑟, and distance to object boundary 𝑏𝑑 [54]. Training follows µSAM’s default configuration, using a Dice loss for each channel together with an additional MSE term:

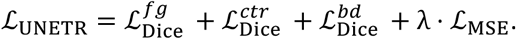

For data split, 60% of images were used for training (1045 cells), 20% for validation (495 cells), and 20% for testing (425 cells). The AIS outputs were evaluated by mean intersection-over-union (IoU), where Full FT ViT-B-LM achieved the highest mean IoU. Thus, we selected this Full FT ViT-B-LM model as YeastSAM.

### Mother-bud separation module

We trained two auxiliary models to automate the mother-bud separation process: (i) a convolutional neural network (CNN) for binary budding/non-budding classification, and (ii) a U-Net for dividing line prediction [46].

Our module operates on binary cell masks rather than brightfield images. This design choice ensures the model’s generality, which makes it possible to apply to other asymmetrically dividing cells beyond just budding yeast. The CNN receives individual mask instances and outputs a class. For each budding cell identified, the U-Net then predicts the dividing line between mother and bud. We found that conventional morphological operations, such as those based on skeletonization or watershedding, were not sufficiently robust for this task, struggling with variations in cell shape and orientation. This motivated our choice of a data-driven U-Net approach.

#### Label generation

To construct training data for dividing line prediction, we first merged the annotated mother–bud masks into a single connected binary mask. Then, we generated dividing line labels by applying a distance-transform–based bridging strategy: the shortest path between the two touching or closely adjacent cell regions was identified, and its centerline was extracted as the dividing line. As the dividing line only accounts for few pixels, the class distribution within each mask is highly imbalanced. To alleviate severe class imbalance, the line labels were dilated to a fixed thickness [55]. Each sample was cropped to the bounding box of the cell pair, padded, and resized to 256×256. These transformations are reversible, which allows to revert to original cell masks during inference.

### Loss function

For the CNN classifier, we adopted a weighted cross-entropy loss to correct for class imbalance. For the U-Net dividing line predictor, dividing line pixels are sparse relative to background, so we adopted a combined loss to balance precision and overlap, including focal loss, Dice loss, and binary cross entropy [56]. The focal loss is defined as:

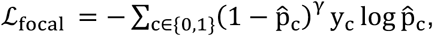

where y_&_ is the ground-truth label, p9_&_ is the predicted probability of class c, and γ is a focusing parameter controlling the strength of down-weighting for easy samples. The final combined loss is formulated as:

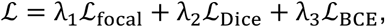

with empirically chosen weights λ_=_, λ_@_, λ_A_.

#### Training setup

The CNN classifier and the U-Net dividing line predictor were trained separately. For the CNN training, we used the Adam optimizer (learning rate 1e-3, weight decay 1e-4), batch size 128, and early stopping after 15 epochs. For the U-Net, training was performed with Adam (learning rate 5e-5), batch size 16, up to 500 epochs, and early stopping with patience of 50 epochs.

#### Post-processing

The predicted junction mask is subtracted from the original cell mask to isolate the two primary cell bodies. The geometric centroids of these resulting components are calculated to define a mother-bud axis. The final dividing line is then constructed perpendicular to this axis and positioned to pass through the centroid of the initial junction prediction.

#### Volume estimation

To estimate cell volumes, we segmented cell masks and approximated the 3D shape using the conical method. The cross-sections were rotated along its major axis to form truncated cones. This approach provided approximate volumes for mother and bud compartments, which were subsequently used to calculate mRNA concentrations.

#### Evaluation

We calculated Detection Rate (DR) to evaluate the AIS outputs. The key advantage of DR is that it directly models the biological task of budding cell recognition. In practice, a model may achieve a high IoU score by accurately segmenting cell shapes, but still fail to recognize a budding event if the mother–bud pair is not correctly identified. DR avoids this issue by explicitly assessing whether budding cells are detected as budding, and non-budding cells as non-budding, which reflects the true biological relevance of the task. We also benchmarked conventional mean IoU for fairness. YeastSAM achieved the highest mean IoU among compared methods, in addition to its superior DR. Thus, the preference for DR in our main evaluation does not indicate a weakness in IoU performance, but rather emphasizes an evaluation metric that better aligns with the biological objectives of budding event detection.

For the budding/non-budding classifier, performance was assessed on a held-out test set using accuracy on budding/non-budding cells (precision and specificity). For the U-Net dividing line predictor, we evaluated trained model using Dice coefficient and mean IoU between prediction and GT dilated dividing lines. In addition, we manually inspected predicted dividing lines to assess whether they produced a reasonable separation of mother and bud cells, since a line can be useful even if it does not perfectly overlap with the ground-truth annotation.

YeastSAM was compared with widely used yeast cell segmentation methods by DR, including µSAM (ViT-L-LM), Cellpose-SAM (CPSAM), Cellpose3, YeastMate and YeaZ, with parameters adjusted according to recommended settings.

## Availability and implementation

Documentation for YeastSAM can be found at https://yeastsamdoc.readthedocs.io/en/latest/. All source code for training models can be found at GitHub: https://github.com/YonghaoZhao722/YeastSAM.

---

**Supplementary Movie 1. Image annotation of YeastSAM.**

The movie illustrates how a mother-bud pair initially misclassified as two G1-phase cells was corrected. First, the incorrect mask was removed. Next, guidance points were added: green points indicating regions inside the mask and a red point indicating a region outside. The program then generated the corrected mask. This procedure was repeated to correct all misclassified cells.

**Supplementary Figure 1.**
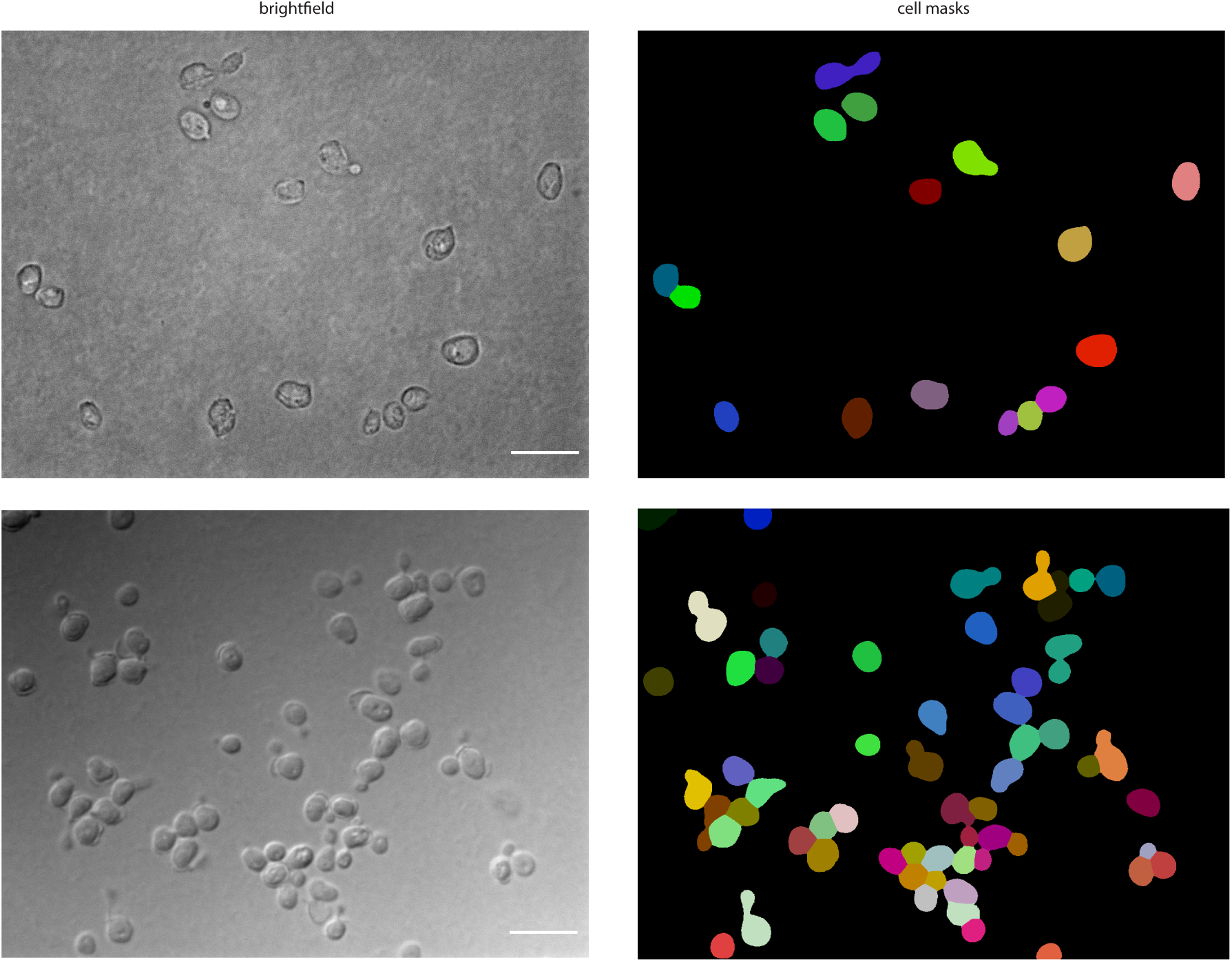
Examples of YeastSAM segmentation. Brightfield images with corresponding YeastSAM segmentation outputs. Individual cells are shown in distinct colors. Scale bars, 10 μm.

**Supplementary Figure 2.**
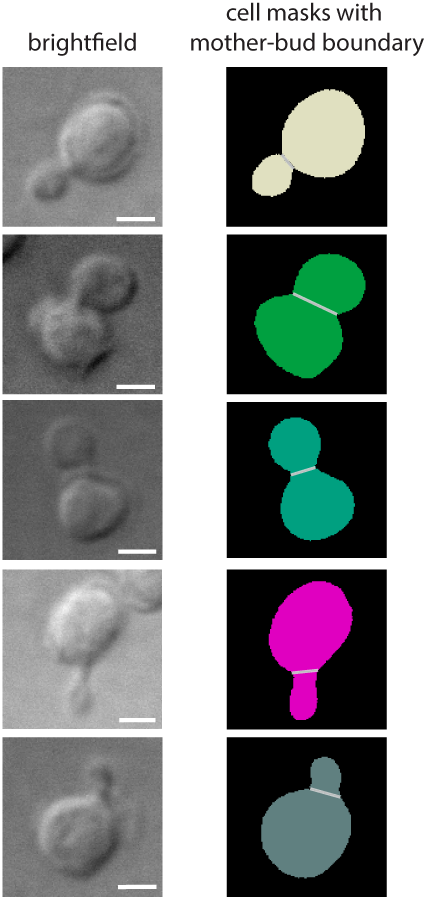
Example cells analyzed with YeastSAM and mother-bud separation module. Brightfield images with corresponding segmentation outputs. Mother-bud boundaries are indicated in grey. Scale bars, 2 μm.

**Supplementary Table 1.**
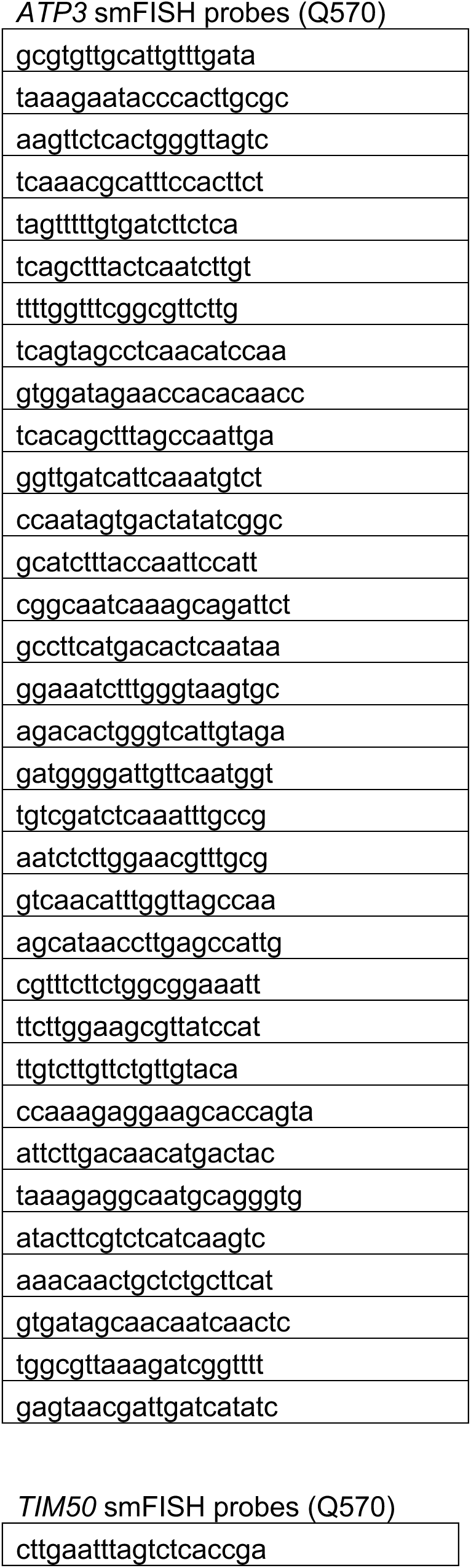
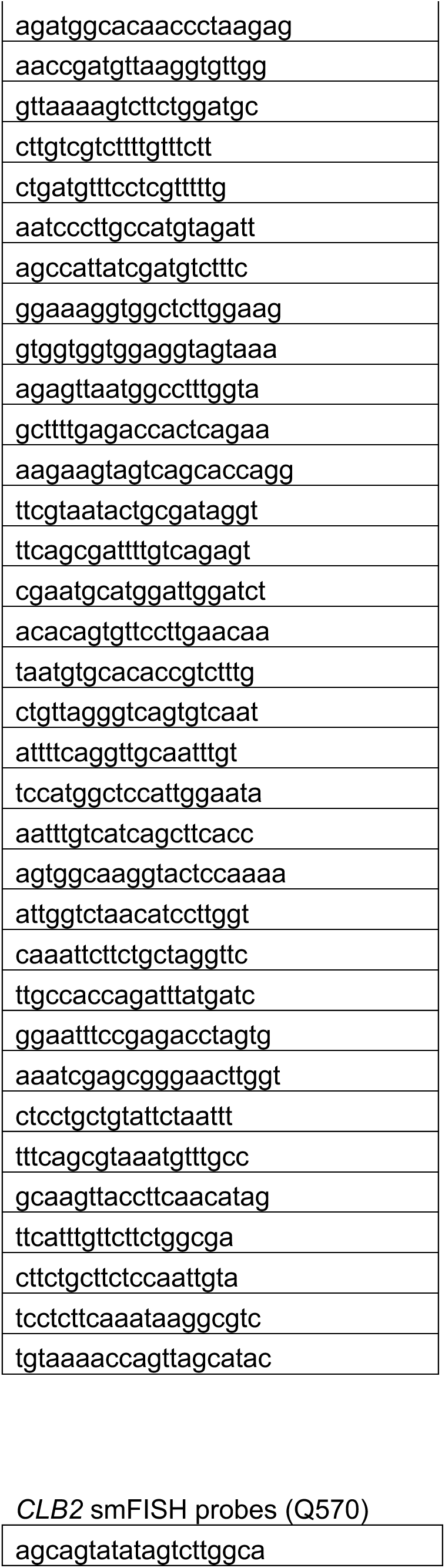
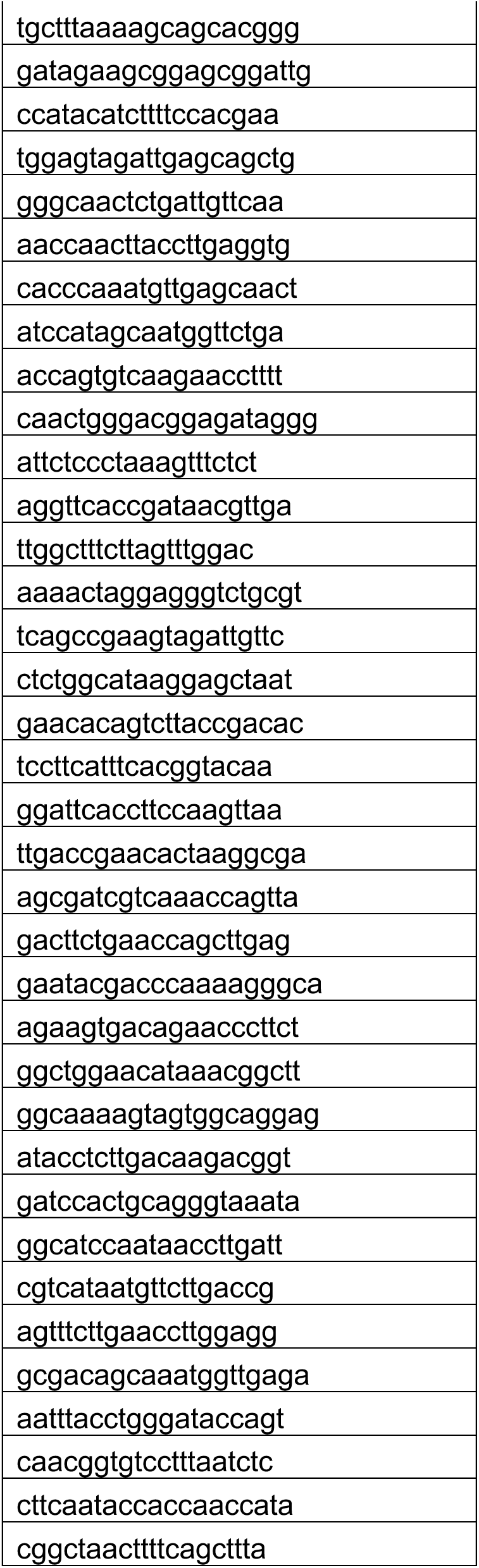
smFISH probes used in this study.

